# Computational Disease Subtyping based on Joint Analysis of Clinical and Genomic Data

**DOI:** 10.1101/307751

**Authors:** Diana Diaz, Aliccia Bollig-Fischer, Alexander Kotov

## Abstract

**Objective:** To investigate application of non-negative tensor decomposition for disease subtype discovery based on joint analysis of clinical and genomic data.

**Data and Methods:** Somatic mutation profiles including 11,996 genes of 503 breast cancer patients from the Cancer Genome Atlas (TCGA) along with 11 clinical variables and markers of these patients were used to construct a binary third-order tensor. CANDECOMP/PARAFAC method was applied to decompose the constructed tensor into rank-one component tensors. Definitions of breast cancer verotypes were constructed from the patient, gene and clinical vectors corresponding to each component tensor. Patient membership proportions in the identified verotypes were utilized in a Cox proportional hazards model to predict their survival.

**Results:** Qualitative evaluation of the verotypes obtained by tensor factorization indicates that they correspond to clinically meaningful breast cancer subtypes. While some components correspond to the known HER2- or ER-positive breast cancer subtypes, other components correspond to a variant of triple negative subtype and a cohort of patients with high mutation load of tumor suppressor genes. Quantitative evaluation indicates that the Cox model utilizing computationally discovered breast cancer verotypes is more accurate (AUC=0.5796) at predicting patient survival than the Cox models utilizing random patient membership proportions in cancer subtypes (AUC=0.4056) as well as patient membership proportions in genotypes (AUC=0.4731) and phenotypes (AUC=0.5047) obtained by non-negative factorization of the somatic mutation and clinical matrices.

**Conclusion:** Non-negative factorization of a binary tensor constructed from clinical and genomic data enables high-throughput discovery of breast cancer verotypes that are effective at predicting patient survival.

## BACKGROUND AND SIGNIFICANCE

Successful transition into the era of precision medicine^1,2^ or screening, diagnostic, therapeutic and prognostic procedures that take into account individual variability of patients, requires comprehensive knowledge of complex relationships between molecular, biological and physiological processes in a human body. Stratification of patients into cohorts with a common biological basis and phenotypic manifestation of a particular disease is an important aspect of such knowledge. Remarkable advances in the next-generation sequencing technology coupled with widespread adoption of electronic health records (EHR) by healthcare providers in the United States have enabled collection of unprecedented amounts of genetic and clinical patient data, from which such knowledge can be discovered. Specifically, methods for high-throughput computational analysis of genetic and clinical data can help shed light on heterogeneous (molecular, biological and physiological) markers that are highly predictive of survival as well as the outcome of therapeutic agents and treatment strategies. Prior research along this direction has focused on the methods to analyze genetic and clinical data in isolation with the goal of identifying either phenotypes (i.e. sets of biomarkers that are more prevalent in individuals with a particular disease or condition than in the general population)^3–9^ or genotypes (i.e. DNA sequences that underlie specific diseases or traits)^10–14^. In particular, the recently proposed computational methods for discovering EHR-based phenotypes have been successfully applied to patient cohort identification^8^ and determining the eligibility of patients for clinical trials.^15^ On the other hand, the mapping of the human genome has enabled computational genotyping methods, which typically combine clustering^10,11,16^ with data integration^12,17^ and feature selection^18,19^ to identify the genes that are predictive of specific clinical outcomes.^20–22^ Previous research on personalized approaches to cancer treatment has primarily focused on genetic studies, including identification of pathogenic mutations of individual genes in cancer tumors,^14^ tumor stratification,^22–25^ functional diagnostics,^26^ and classification,^27^ as well as creating centralized resources, repositories and protocols for interpreting, validating, sharing and updating the results produced by these studies.^28^

Despite the immense progress in computational methods for analyzing clinical or genomic data, neither of these two types of methods alone can capture all aspects of the pathogenesis of complex diseases,^29^ such as cancer. The emergence of EHR-linked biobanks, such as those created by the Electronic Medical Records and Genomics (eMERGE) consortium,^30^ enabled computational methods to discover associations between specific diseases and genes^31 -34^ through genome-wide association studies (GWAS)^35^ or between specific phenotypes and genes through phenome-wide association studies (PheWAS).^36^,^37^

This paper reports the results of applying computational methods for high-throughput cancer subtyping based on the joint analysis of clinical and genomic data of breast cancer patients and utilizing the discovered subtypes for their survival prediction. Breast cancer is the most frequently diagnosed cancer and the second cause of cancer deaths in women.^38^ Due to a significant degree of variation, cancers are particularly difficult to treat, since patients with the same type of cancer and a similar set of symptoms can have different responses to the same treatment.^39,40^ One in three women with breast cancer detected by a mammogram unnecessarily undergo surgery or chemotherapy,^41^ which results in measurable toll of increased healthcare costs and unmeasurable emotional trauma. Therefore, methods for more comprehensive and accurate stratification of cancer patients would help oncology decisionmaking process, facilitate treatment selection and ultimately improve patient outcomes. There is a particular need for such methods in the case of women with early stages of breast cancer, since there is a great variety in treatments prescribed to those women, with some women being under- and over-treated, as a result.

Since cancer development and progression are influenced by several factors, including germ-line or somatic tumor genetics, overall patient health as well as environmental or lifestyle factors,^42^ it is natural to assume that cancer subtypes should incorporate all these different modalities of patient data. However, existing integrative approaches are specific to one particular type of clinical data, such as chemistry evaluations,^43^ survival,^20,44^ or epidemiological data,^45^ and there has been relatively little research on computational methods for joint analysis of clinical and genomic data for disease subtyping.

To address this deficiency, we propose CLIGEN, a high-throughput pipeline for fully unsupervised disease subtyping based on CLInical and GENomic data. Specifically, we focus on the problem of identifying cohorts of patients, which share the same set of pathogenic gene mutations as well as the same values of clinical variables and markers. This problem is different from tumor stratification, which is based only on genetic information and is aimed at dividing the heterogeneous population of cancer tumors into biologically meaningful subtypes based on mRNA expression data^22,24^ or gene networks.^23^ Although genome-scale molecular information provides an insight into biological processes driving tumor progression, cancer subtyping based on gene expression profiles alone has been shown to have limited correlation with clinical outcomes.^46,47^

In the first stage of the proposed pipeline, multi-modal patient data that includes somatic mutation profiles as well as clinical variables and markers is represented as a binary three-dimensional tensor. As differential measurements between a tumor and normal tissue, somatic mutation profiles are more suitable for disease subtyping than other types of ‘omics’ data, which are absolute measurements for each patient. Furthermore, somatic mutations capture causal genetic events underlying tumor progression, whereas mRNA or protein expression profiles are functional readouts of the current cell state and can be influenced by external factors that are unrelated to tumor biology.

In the second stage of the proposed pipeline, non-negative tensor decomposition is applied to identify latent factors in each modality of the constructed tensor. These latent factors correspond to the frequently cooccurring combinations of gene mutations and clinical markers in patients with a particular complex disease, such as cancer. We hypothesize that the proposed pipeline enables discovery of cancer verotypes (clinical disease subtypes combining phenotypes and genotypes).^48^ To validate this hypothesis, we applied the proposed framework to discover breast cancer verotypes based on the clinical and genetic data in the Cancer Genome Atlas (TCGA)^49^ and experimentally demonstrate that the discovered breast cancer verotypes can provide actionable insights, such as patient survival prognosis, to clinicians at the point of care.

## DATA AND METHODS

### Dataset description

We used somatic mutation (non-silent mutation from the whole exome sequencing level 3) profiles and clinical data of breast cancer patients from The Cancer Genome Atlas (TCGA)^49^ for all experiments in this work (data downloaded from cBioportal on 9 April 2017).^50^ We considered only the patients for whom both somatic mutation and clinical data were available and discarded the patients with fewer than 10 somatic mutations. This resulted in a dataset combining information about mutations in 11,996 genes and 70 dichotomized values of 11 clinical variables and markers for 503 patients.

#### Somatic mutations

The somatic mutation table consists of 37 columns and 34032 registries. A registry in this table indicates a mutation in the gene reported in the column *‘Hugo Symbol’* for the sample in the column *‘Tumor Sample Barcode’*. Additional details on processing and organization of these data are available in.^50^ In this work, we constructed patient mutation profiles as binary vectors, in which a bit is set, if the patient’s gene corresponding to that position in the vector harbors a mutation.

#### Clinical variables and markers

The clinical variables and markers (without loss of generality, further referred to as clinical markers) in TCGA include the age of diagnosis with cancer, gender, estrogen receptor (ER) status, progesterone receptor (PR) status, human epidermal growth factor receptor 2 (HER2) final status, American Joint Committee on Cancer (AJCC) breast cancer stage, AJCC coded tumor stage (T), AJCC coded lymph node stage (N), AJCC coded metastasis stage (M), IHC (immunohistochemistry) expression level and PAM50 profile.

### Proposed pipeline

CLIGEN, the proposed pipeline for unsupervised verotyping of complex diseases, is illustrated in Figure 1. The input to the pipeline consists of genetic and clinical data of the patients with the same complex disease (e.g. cancer). The output of the pipeline is a set of definitions of disease subtypes characterized by genetic and clinical markers. The pipeline consists of the two stages: 1) data preprocessing and tensor construction and 2) non-negative decomposition of the constructed tensor to obtain cancer subtypes. Detailed descriptions of each stage are provided below and the notations used in these descriptions are summarized in Table 1. Code for CLIGEN is available at http://www.cs.wayne.edu/dmd/cligen/.

**Figure 1.**
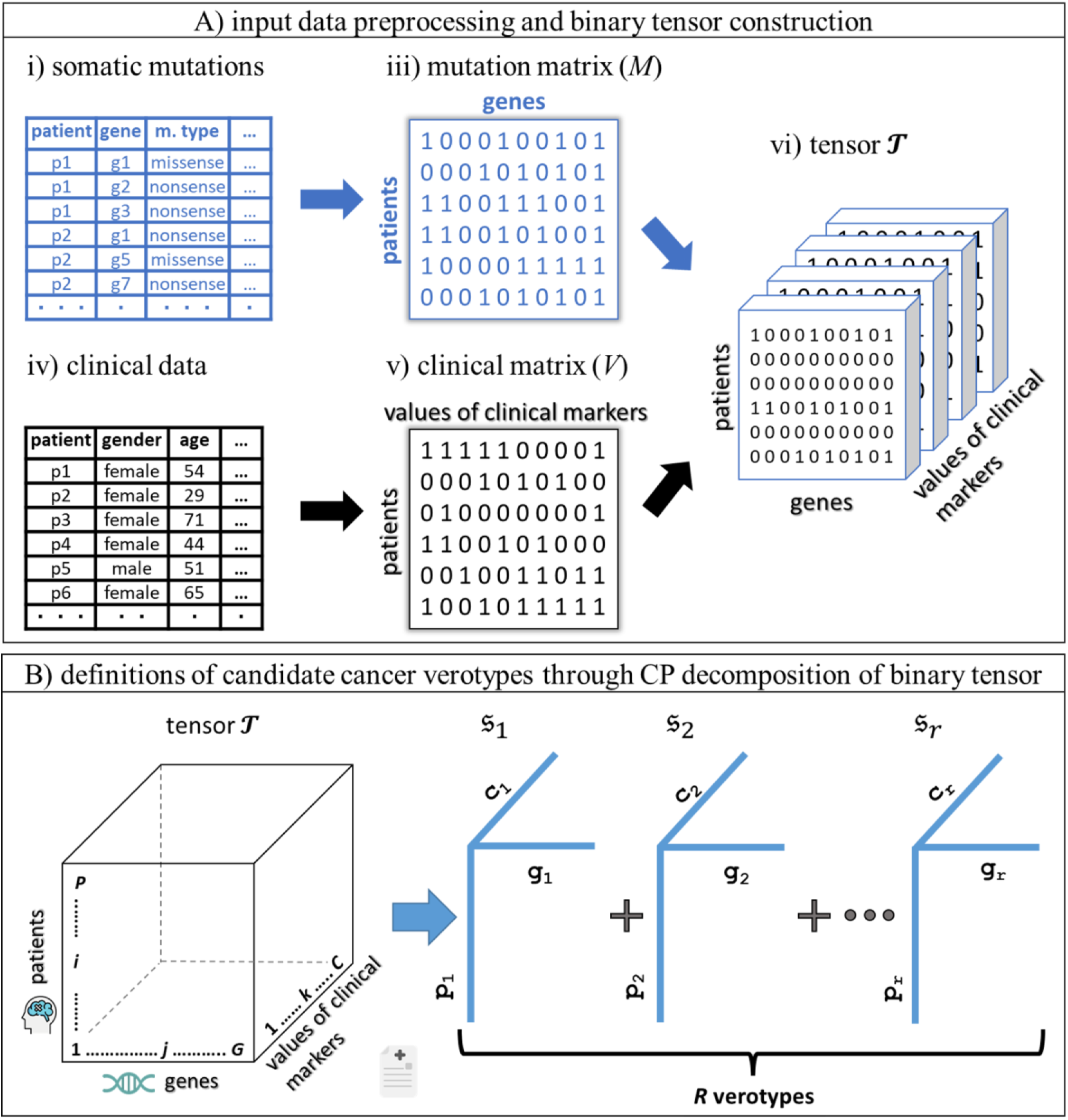
Two stages of the proposed CLIGEN pipeline: A) data preprocessing and construction of multi-modal three-dimensional tensor 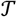. B) obtaining candidate verotypes via CP decomposition of tensor 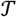.

**Table 1.** List of notations used in this paper and their definitions

| Symbol | Definition |
| --- | --- |
| $M, V$ | somatic mutation and clinical matrices |
| $M_{ij}$ | cell of mutation matrix for patient $i$ and gene $j$ |
| $V_{ik}$ | cell of clinical matrix for patient $i$ and $k$-th value or interval of clinical markers |
| $\mathcal{T}$ | multi-modal binary CLIGEN tensor |
| $P, G, C$ | number of patients, genes as well as intervals and values of clinical markers |
| $i, j, k$ | indices of patients, genes, and clinical markers. |
| $t_{ijk}$ | value of tensor cell for the $i$-th patient, $j$-th gene, and $k$-th value or interval of clinical markers |
| $\circ$ | outer product of two vectors |
| $R$ | number of tensor components |
| $s_r$ | $r$-th rank-one component i.e. a candidate verotype definition |
| $p_r, g_r, c_r$ | patient, gene and clinical factor vectors |

### Data preprocessing

The first stage of the proposed pipeline, illustrated in Figure 1A, involves preprocessing the input genomic and clinical patient data, to create a combined representation for subsequent analysis.

#### Somatic mutation data

Somatic mutation datasets typically take the form of mutation tables, in which the rows correspond to mutations and the columns describe the type of each mutation and its location (Figure 1Ai). Based on the input mutation table, CLIGEN constructs a binary mutation matrix *M* with patients as rows and genes as columns. The value of 1 in the cell *M_ij_* of matrix *M* indicates that patient *i* has at least one non-silent mutation (i.e., a mutation of any of the following types: missense mutation, nonsense mutation, non-stop mutation, in-frame insertion, in-frame deletion, or frameshift mutation) in gene *j*, while the value of 0 indicates that patient *i* has no mutations in gene *j*.

#### Clinical data

Continuous clinical variables or markers, such as the age of diagnosis, were discretized into intervals with a total of *n* combined intervals of all continuous markers and levels of all discrete markers in the entire dataset. The values of clinical markers for each patient were represented as abinary clinical matrix *V* with patients as rows and intervals of continuous or levels of discrete clinical markers as columns. The value of 1 in the cell *V_ik_* of matrix *V* indicates that patient *i* has a *k*th level or interval of a discrete or continuous clinical marker.

### Tensor construction

Clinical and mutation matrices are combined to create a threedimensional binary CLIGEN tensor (i.e. multidimensional array) 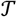 representing interactions between clinical markers and genetic mutations. The first dimension (i.e. mode) of tensor 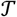 corresponds to patient data, while the other two dimensions correspond to the values of clinical markers and genes (Figure 1B). Each element *t_ijk_* of the tensor 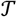 has the value of 1, if the *i*th patient has at least one mutation in gene *j* and *k*th value of clinical markers and 0 otherwise.

Slicing tensor 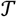 along each of its three modes yields the following views:

1. *Patient mode:* each slice is a matrix of co-occurrences of mutations and clinical markers for a particular patient. F or instance, if a patient’s health records indicate stage I breast cancer and her mutation profile indicates a mutation in gene TP53, then a cell at the row for the “Stage I” clinical variable and the column for the TP53 gene in the matrix corresponding to the tensor slice for this patient will have a value of 1. The cells in the same column and the rows for “Stage II”, “Stage III” and “Stage IV” will have the value of 0.
2. *Gene mode:* each slice is a matrix with patients as rows and clinical markers as columns, which shows how a mutation in a particular gene is correlated with clinical markers in different patients. Such matrix can be considered as a summary of phenotypic manifestations of a particular gene mutation.
3. *Clinical mode:* each slice is a matrix with patients as rows and genes as columns, which shows how gene mutations in different patients are correlated with a particular clinical marker. Such matrix can be considered as a summary of genetic markers for a simple phenotype.

#### Obtaining candidate verotype definitions through tensor factorization

Tensor decomposition^51^ is a powerful mathematical technique that has been successfully applied in different domains ranging from psychology and neuroscience to computer vision.^52^ In biomedical informatics, tensor decomposition has proven to be useful for understanding cellular states^53^ and biological processes,^54^ in addition to EHR-based phenotyping^3–5^. Tensor decomposition has several advantages over matrix factorization. First, tensors explicitly account for the multiway structure of the data that is otherwise lost, when a tensor is converted into a matrix by collapsing some of its modes. Second, some tensor decomposition methods guarantee uniqueness of the optimal solution even for very sparse tensors. The two most widely used tensor decomposition methods are the Tucker method^55^ and CP, which stands for Canonical Decomposition (CANDECOMP),^56^ also known as Parallel Factor Analysis (PARAFAC).^57^ CP decomposes a tensor into a linear combination of rank-one tensor components.^56^ Each component of the CLIGEN binary tensor consists of vectors corresponding to each mode,^51^ with each vector corresponding to groups of frequently co-occurring genes, clinical markers or patients. The CP decomposition can be considered as a special case of the Tucker decomposition when the size of each modality of the core array is the same and the only non-zero elements in the core are the elements along the main diagonal.^51^ An important property of CP decomposition is that the restriction imposed on the Tucker core leads to the uniqueness of the optimal solution.^51^

Specifically, CP decomposition of a third-order CLIGEN tensor 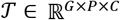 with dimensions corresponding to genes, patients, and clinical markers is formally defined as:

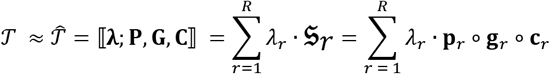

where *R* is the number of component rank-one tensors 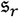 that tensor 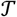 is decomposed into and 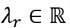 is the weight of the *r*th component. Each co mponent is associated with three length normalized non-negative vectors: a patient vector 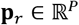, a gene vector 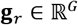 and a clinical vector 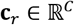. Patient, gene and clinical vectors for all tensor components correspond to the columns of the patient, gene and clinical factor matrices **P, G, C**.

CP decomposition is obtained by solving the following optimization problem:

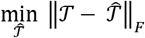

aimed at finding the best approximation 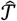 of each element *t_ijk_* of the original tensor 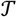 from the factor vectors corresponding to the components tensors as follows:

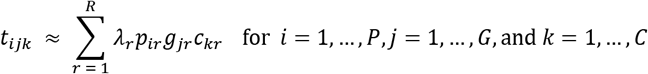

Verotype definitions are constructed from patient, gene and clinical factor vectors corresponding to each of the component tensors obtained by CP decomposition of the CLIGEN tensor. Each element in the vectors corresponding to gene and clinical tensor modes can be interpreted as an importance of a gene or clinical marker to a verotype. The prevalence of a verotype among patients is calculated as a proportion of non-zero entries in the patient vector corresponding to a given verotype.

## RESULTS

We performed both qualitative and quantitative evaluation of breast cancer subtypes derived from TCGA breast cancer dataset^50^ using the proposed pipeline.

### Quantitative evaluation

Quantitative evaluation was conducted for the task of patient survival prognosis, which is important for personalizing cancer treatment.^58^ Specifically, we compared Cox proportional hazards models that use the following predictors for survival prognosis of breast cancer patients:

M1: membership proportions of a patient in breast cancer verotypes, which correspond to a row in the patient factor matrix **P** obtained by the proposed pipeline;
M2: membership proportions of a patient in breast cancer genotypes, which correspond to a row in matrix **P** obtained by nonnegative factorization of the somatic mutation matrix *M* (as in [23] but without gene network smoothing), as shown in Figure 2;
M3: membership proportions of a patient in breast cancer phenotypes, which correspond to a row in the patient factor matrix obtained by non-negative factorization of the clinical matrix *V*;
M4: random membership proportions of a patient in each number of breast cancer subtypes

In the first experiment, we compared the accuracy of the Cox models using each of the above predictors for survival prognosis of breast cancer patients, while in the second experiment, we compared the goodness of fit of these models.

**Figure 2.**
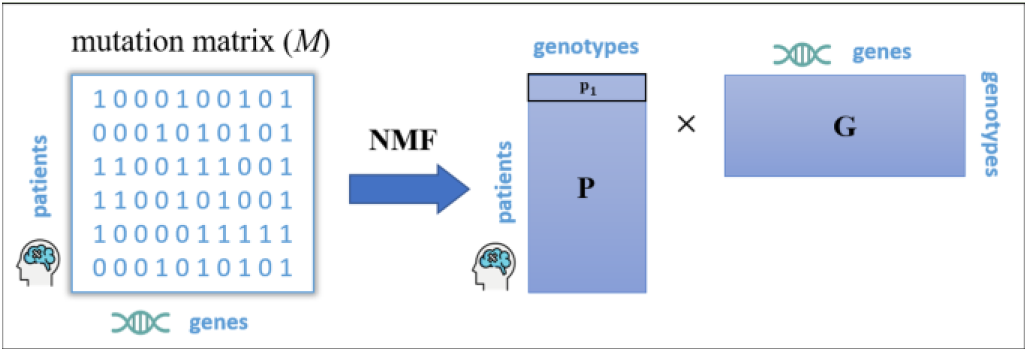
Obtaining genotypes through non-negative factorization of a binary somatic mutation matrix. The rows in matrix P correspond to a vector of genotype memberships (e.g. p_1_ is a vector of genotype memberships for the first patient). Rows in matrix G correspond to genotype definitions.

#### Accuracy of survival prognosis

In the first experiment, we compared the area under the ROC curve (AUC) for the models M1-M4. We used 10 random splits, such that each split divides the TCGA patient data into two halves (50% for testing and 50% for training), as experimental design. The Cox models were estimated using the data in the training splits and evaluated using the data in the testing splits.

The plot of AUC values for models M1-M3 micro-averaged over splits by varying the number of the most prevalent cancer verotypes, genotypes and phenotypes is shown in Figure 3, from which two major conclusions can be drawn. First, the Cox regression model that utilizes patient membership proportions in verotypes obtained by CLIGEN (M1) is consistently more accurate at predicting breast cancer survival than the Cox model that uses membership proportions in genotypes obtained by NMF (M2) and phenotypes obtained by NMF (M3), which indicates the importance of taking into account both clinical and genomic data when determining cancer subtypes. In particular, the Cox model utilizing patient verotype membership proportions as predictors achieved the highest AUC of 0.5796, when 10 most prevalent verotypes were used, while the Cox model utilizing patient genotype memberships as predictors achieved the highest AUC of 0.4731, when the 9 most prevalent genotypes were used, and the Cox model utilizing patient phenotype memberships as predictors achieved the highest AUC of 0.5047, when the 6 most prevalent phenotypes were used.

**Figure 3.**
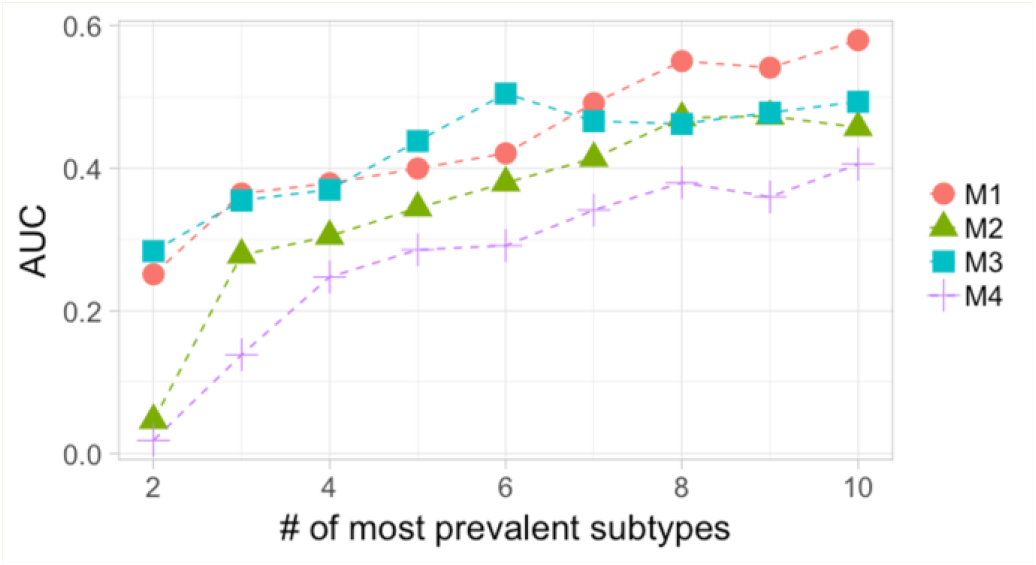
AUC of Cox models for breast cancer patient survival prediction that utilize patient membership proportions in most prevalent verotypes obtained by the proposed pipeline (M1), patient membership proportions in genotypes (M2) obtained by NMF of binary somatic mutation matrix, phenotypes (M3) obtained by NMF of binary clinical matrix as predictors, and random patient membership proportions (M4).

Second, the Cox models utilizing patient membership proportions in the top-*k* most prevalent verotypes derived by CLIGEN as well as the phenotypes and genotypes derived by NMF are all more accurate at predicting breast cancer survival than the baseline Cox model utilizing random patient membership proportions in the same number of cancer subtypes (AUC = 0.4056).

#### Goodness of fit

In the second experiment, we compared the goodness of fit of the models M1-M4 estimated on the entire TCGA dataset. The p-values of Log-rank and Wald tests of these models are summarized in Table 2. Both tests indicate that patient membership proportions in verotypes derived by CLIGEN are more statistically significant predictors of breast cancer patient survival than membership proportions in breast cancer phenotypes, which in turn are more statistically significant predictors than random patient membership proportions and membership proportions in genotypes derived by NMF.

**Table 2.** p-values of Log-rank and Wald tests of the Cox proportional hazard models utilizing patient membership proportions in verotypes (M1), genotypes (M2), phenotypes (M3), and random membership assignment (M4) as predictors.

| Model | Log-rank test | Wald test |
| --- | --- | --- |
| M1 | 8.327e-15 | 0.000082 |
| M2 | 0.315 | 0.3772 |
| M3 | 0.0007 | 0.0124 |
| M4 | 1.29e-5 | 0.00012 |

Kaplan-Meier survival plots for the 4 most prevalent breast cancer subtypes obtained using CLIGEN and NMF are shown in Figure 4. As follows from Figure 4, the most prevalent subtypes obtained using the proposed pipeline are more distinct in terms of patient survival dynamics (*p* = 0.0493) than the most prevalent subtypes obtained using genotypes (*p* = 0.241) and phenotypes (*p* = 0.2073).

**Figure 4.**
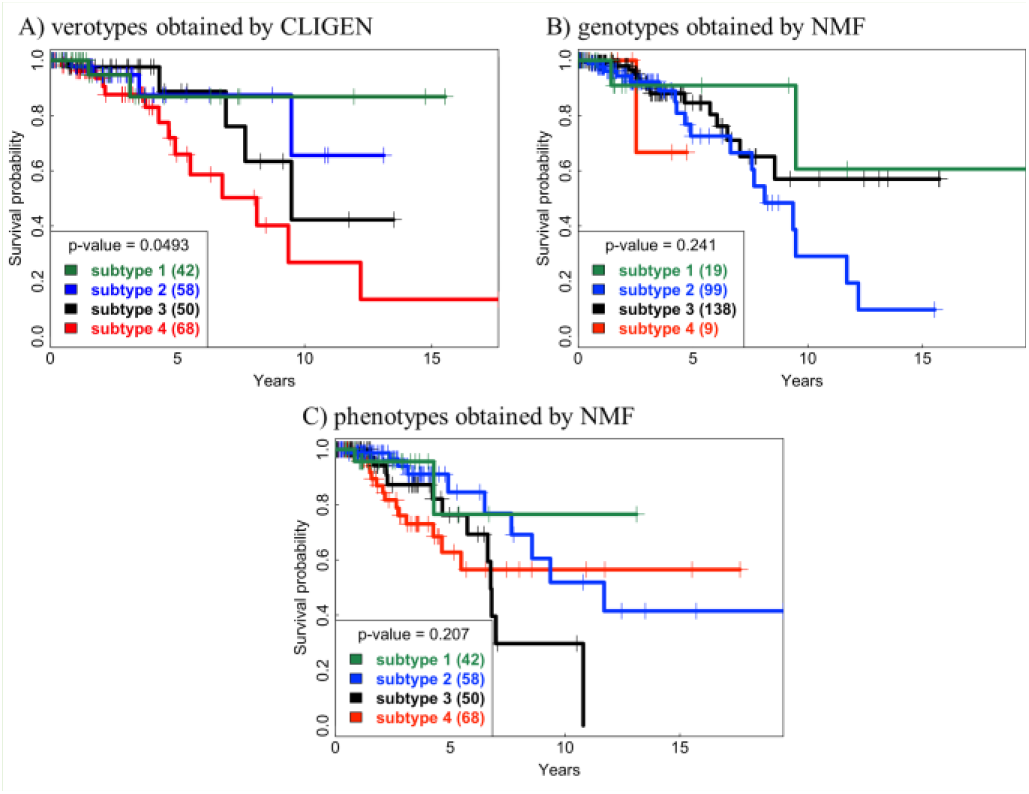
Kaplan-Meier survival plots for the 4 most prevalent: A) verotypes obtained using CLIGEN, B) genotypes obtained using NMF, C) phenotypes obtained using NMF.

### Qualitative evaluation

An oncologist performed qualitative evaluation of breast cancer verotypes obtained by CP decomposition of the binary CLIGEN tensor constructed from the TCGA dataset into 10 components, since this decomposition results in the most accurate prediction of cancer survival. An enrichment analysis of the list of genes associated with each verotype was performed using the Ingenuity Systems Upstream Analysis tool.^59^

#### Examination of verotypes

Each breast cancer verotype obtained by CLIGEN was analyzed and compared with the known breast cancer subtypes. Component 1 (shown in Figure 5A) corresponds to a small cohort of patients with a high mutation load. Further investigation of this verotype revealed a large number of mutations in the tumor suppressor genes (BRCA1, BRCA2, TP53, PTEN, RB1) that participate in DNA repair, which indicates that the high mutation load may be associated with a mutation in a DNA repair gene pathway(s). As follows from Figure 6, for each sample, these mutations were mutually exclusive. Further investigation of these genes can elucidate biological process(es) underlying these mutations.

**Figure 5.**
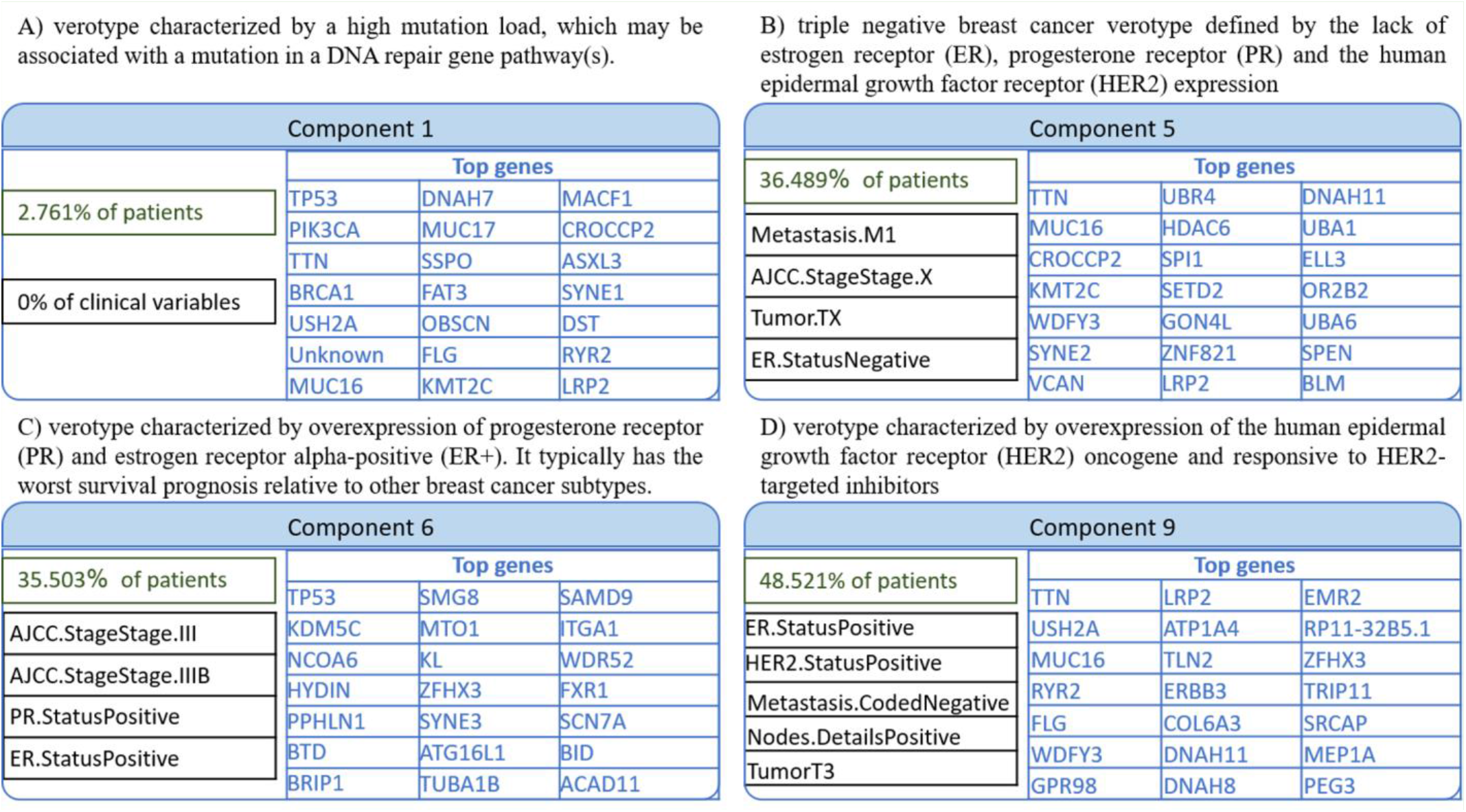
Examples of verotypes identified by the proposed pipeline.

**Figure 6.**
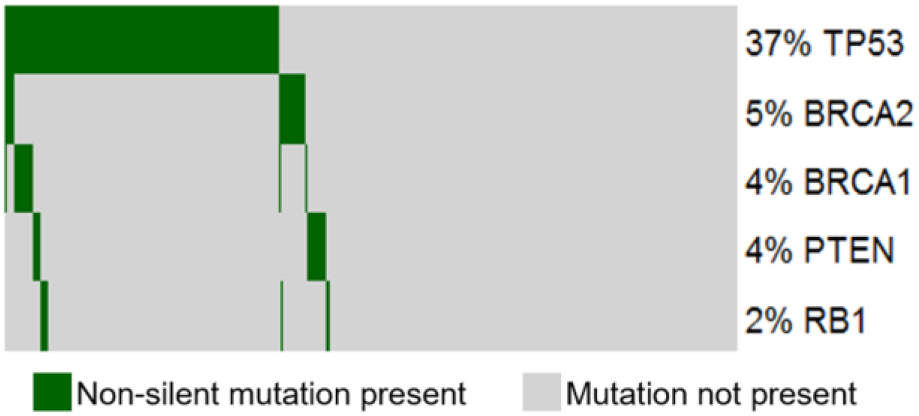
Mutual exclusivity across the characteristic genes of one of the verotypes identified by the proposed pipeline.

Component 5 (shown in Figure 5B) appears to correspond to a subtype of the triple negative breast cancer (TNBC), which is a defined by the lack of ER, PR and HER2 expression. Molecular aberrations driving this breast cancer subtype remain undefined and patients with this subtype of breast cancer have the worst prognosis relative to the patients with any other known breast cancer subtype. Component 6 (shown in Figure 5C) corresponds to the subtype of progesterone receptor and estrogen receptor alpha-positive (ER+) cancers, that are responsive to anti-ER therapies. Based on the associated clinical markers, Component 9 (shown in Figure 5D) appears to be related to the known subtype of breast cancer that is driven by overexpression of the epidermal growth factor receptor oncogene (HER2) and responsive to HER2-targeted inhibitors.

#### Triple negative Breast Cancer

Further analysis of putative cancer driver genes, potentially activated or inactivated by mutations associated with the TNBC-related fifth component resulted in significant enrichment of genes with a role in signaling networks that promote the function of cancer stem-like cells (CSCs), i.e., downstream of transcription factor TWIST1, and alternative mRNA splicing, i.e., downstream of serine and arginine-rich splicing factor SRSF2. CSCs are identified in patient TNBC tumors as a fraction of self-renewing, tumor-initiating cancer cells that also give rise to drug resistance and metastatic recurrence.^60–62^ Alternative mRNA splicing has also been implicated in maintaining and generating CSCs.^63^

## DISCUSSION

Methods for integrative analysis of genomic and clinical data face a common challenge of dealing with large volumes of data. By utilizing sparse representations and inexpensive linear algebra operations, tensor factorization methods effectively address this challenge. Successful application of tensor decomposition in different domains led to further research into efficient optimization methods for tensor decomposition,^64^ which makes tensor decomposition a method of choice for high-throughput disease subtyping.

Since tensor factorization models are parametric, selecting the optimal number of components for CP decomposition of the binary tensor (i.e. model order estimation) is an important practical aspect of the proposed pipeline. Too few components typically result in general verotype definitions, which may combine several actual disease subtypes. Too many components typically result in specific verotype definitions, which may split the actual disease subtypes. It is important to point out that, in terms of the number of model parameters, CP decomposition, which assumes that the number of components is the same per each tensor mode, has an advantage over Tucker decomposition, which requires specifying the number of components per each mode. While it is known that the number of components that minimizes the reconstruction error of the original tensor from its components is equal to the rank of a given tensor,^51,65^ finding tensor rank is an NP-complete problem.^66^ Even if the rank of a tensor is known, the number of components that minimizesreconstruction error may not result in the best accuracy for a particular task, such as survival prediction. Therefore, the optimal number of components is typically determined using heuristics, such as core consistency diagnostic,^67^ cross validation^68^ (as was done in this work) or hierarchical Bayesian approach,^69^ if a suitable prior can be defined.

Tensor construction is another aspect of the proposed approach with possible variations. In this work, we used the presence or absence of non-silent gene mutation as a single genetic signature of patients. However, it is possible to use other types of genomic data, such as gene expression or copy number variation. It is also possible to construct a count tensor, instead of a binary tensor, by taking into account both the type of mutations and the number of mutations per gene, which we leave for future work.

## CONCLUSION

In this paper, we introduced CLIGEN, a novel machine learning based pipeline for unsupervised disease subtype discovery based on non-negative decomposition of a binary tensor combining clinical and somatic mutation patient data, and applied the proposed pipeline to breast cancer subtyping. Qualitative and quantitative evaluation of the discovered breast cancer subtypes indicates that representation of clinical and genomic patient data as a binary tensor and its subsequent nonnegative decomposition is an effective computational approach to high-throughput disease subtyping for precision medicine. In particular, our proposed pipeline was not only able to identify known breast cancer subtypes (HER2+ and ER+), but also elucidated new possible characteristics of a complex breast cancer subtype (triple negative), which provides an opportunity for further research to define new cancer subtypes. We also demonstrated that patient membership proportions in the discovered breast cancer verotypes are more effective predictors of breast cancer survival than patient membership proportions in computationally identified genotypes and phenotypes.

Although the proposed pipeline was evaluated using the data of breast cancer patients, it can, in principle, be applied to computational subtyping of other cancers and complex diseases.

## Acknowledgments

Results presented in this paper are based on the data in The Cancer Genome Atlas managed by the NCI and NHGRI. We would like to thank Dr. Sorin Drăghici for the helpful suggestions regarding this manuscript.

